# Sealing the deal – Antarctic fur seals’ active hunting tactics to capture small evasive prey revealed by miniature sonar tags

**DOI:** 10.1101/2023.10.19.563066

**Authors:** Mathilde Chevallay, Christophe Guinet, Didier Goulet-Tran, Tiphaine Jeanniard du Dot

## Abstract

Fine-scale interactions between predators and their prey are key factors determining both predators’ hunting efficiency and prey survival. The ability to adopt hunting tactics that minimise the risk of triggering an escape reaction from the prey is crucial for an efficient foraging, and is contingent to detection capabilities and locomotor performances of both predators and prey. In this study, we aimed at describing fine-scale predator-prey interactions in female Antarctic fur seals (*Arctocephalus gazella,* AFS hereafter), a small pinniped foraging on evasive prey. Our objectives were (1) to describe characteristics of prey targeted by female AFS to assess prey selection in this species; (2) to estimate the timing of prey detection by AFS and prey reaction to the AFS approach, and (3) to describe AFS hunting tactics, i.e. fine-scale AFS posture and swimming activity during prey capture events. *In fine*, we seek to better understand AFS’s efficiency at exploiting their small prey. To do so, we used data recorded by a newly developed sonar tag that combines active acoustics with ultra-high resolution movement sensors to study simultaneously the fine-scale behaviour of both AFS and prey during predator-prey interactions. We analysed more than 1200 prey capture events in eight female AFS and showed that AFS and their prey detect each other at the same time, i.e. 1-2 seconds before the strike, forcing AFS to display reactive fast-moving chases to capture their prey. Prey detection was consistently followed by bursts of accelerations from AFS, which lasted longer for evasive prey. We suggest that the ability of AFS to perform bursts of accelerations may allow them to target evasive prey. This active hunting tactics is likely very energy-consuming but might be compensated by the consumption of highly nutritious prey.

## Introduction

Fine-scale predator-prey interactions, i.e. how predators find, select and capture their prey, and alternatively how prey detect and react to imminent predation, are critical in determining both predators’ hunting efficiency and prey survival (Cooper Jr, 1997; McHenry et al., 2009; Stewart et al., 2013), and in shaping their population dynamics (Estes & Duggins, 1995; Frederiksen et al., 2006; Letnic et al., 2012). Sensory capabilities and locomotor performances, i.e. manoeuvrability and acceleration abilities, of both predators and prey are key factors in determining the outcome of predator-prey interactions (Domenici & Blake, 1997).

In the three-dimensional marine environment, prey can escape in every direction (Domenici & Blake, 1997), forcing predators to perform energy-expensive rapid manoeuvres and bursts of accelerations. Predators are often larger than their prey, and manoeuvrability being inversely proportional to body size (Henriques et al., 2021; Tucker & Rogers, 2014), prey usually benefit from a greater manoeuvrability (Domenici, 2001). Therefore, large vertebrate predators may be at disadvantage when capturing small elusive prey. By adopting a stealthy approach, predators can delay or even prevent prey flight reaction, avoiding energy-expensive chase (Cooper Jr, 1997, 2003; Meager et al., 2006; Webb, 1984) (Chevallay et al., 2023). However, this stalking tactic implies that predators detect their prey far enough that they have time to adapt their approach tactics before prey detect them (Chevallay et al., 2023; Snyder et al., 2007; Vance et al., 2021). Conversely, if predators and prey detect each other at the same time, predators must adopt a more reactive approach, with quick reactions to catch alerted prey (Snyder et al., 2007). This reactive hunting mode requires predators to be able to perform fast-start manoeuvres.

In the aquatic environment, marine predators rely on different sensorial systems to locate their prey. Echolocating toothed whales can detect prey at long ranges (Jensen et al., 2018; Tønnesen et al., 2020), while non-echolocating predators such as pinnipeds must rely on vision or tactile cues (Adachi et al., 2022; Dehnhardt et al., 1998; Levenson & Schusterman, 1999; McGovern et al., 2015) that may be only detected at short ranges in the dark ocean. While forward motion of large predators creates a bow wave that can be detected at distance by their prey (Blaxter & Fuiman, 1990; McHenry et al., 2009; Stewart et al., 2013), prey can easily go cryptic for non-echolocating predators by staying motionless. This implies that, in most cases, prey should detect their predators before being detected by them. Nevertheless, a recent study highlighted unexpected detection abilities in Southern elephant seals *(Mirounga leonina)*, allowing seals to set up stealth approach tactics (Chevallay et al., 2023). Prey detection capacities of free-ranging marine predators are however poorly studied and it is not clear if other pinniped species benefit from the same capacities as elephant seals.

Antarctic fur seals *(Arctocephalus gazella,* AFS hereafter) are small pinnipeds foraging on mesopelagic prey, mostly myctophids (Jeanniard-du-Dot, Trites, et al., 2017; Lea, Cherel, et al., 2002). As air-breathing diving predators, they must regularly return to the surface to breathe, limiting their time spent at the bottom of dives searching and hunting for prey. Although foraging locations and diet of AFS are well known, fine-scale characteristics of their prey (i.e. prey size and behaviour) and their hunting tactics to subdue their prey are still poorly known. Compared to hind-flipper thrusting and low-expenditure swimming of elephant seals (Burkhardt & Frey, 2008), fast-swimming AFS rely on their fore-flippers for propulsion (English, 1976). They are highly manoeuvrable and able to perform fast bursts of accelerations. We thus hypothesize that AFS rely on these abilities to target evasive, more reactive and faster moving prey than the slow moving and less manoeuvrable elephant seal, which in turn likely relies on highly sensitive sensory systems to surprise their prey (stalking approach, (Chevallay et al., 2023)). The fact that female AFS from Kerguelen tend to forage mainly on the fast-swimming lipid-rich *Gymnoscopelus sp*. myctophid species (Jeanniard-du-Dot, Trites, et al., 2017; Lea, Cherel, et al., 2002; Lea et al., 2008) capable of avoiding trawl net (Guinet et al., 2001) supports our hypothesis that they rely on active-hunting tactics to target large calorific prey.

To better understand prey selection and hunting tactics in AFS and test our hypothesis, we took advantage of a newly developed miniature sonar and movement tag that allow recording fine-scale information of prey characteristics simultaneously to fine-scale AFS behaviour. Our objectives were (1) to describe characteristics of prey targeted by female AFS to assess prey selection in this species; (2) to estimate the timing of prey detection by AFS and prey reaction to the AFS approach, and (3) to describe AFS hunting tactics, i.e. fine-scale AFS posture and swimming activity during prey capture events to better understand how AFS efficiently exploit their small prey.

## Material and methods

### Device deployments and data collection

Data were collected on eight lactating female AFS in December 2022 at Pointe Suzanne, Kerguelen Island (49°26′S–70°26′E, Southern Ocean) under the ethical regulation approval of the French Ethical Committee for Animal Experimentations (#37480-2022052514544991v7) and the Committee for Polar Environment (A2021-48). Females were captured with a hoop net, anesthetised with isoflurane gas, measured (± 1cm) and weighted (± 10g). They were either equipped with a head-mounted DTAG-4 sonar tag (n = 5, 85 x 45 x 20 mm, 120 g in air, see Goulet et al. (2019) for further details) or a head-mounted DTAG-4 mini sound tag (n = 3, 68.0 x 31.0 x 20.7 mm, 58 g in air). Tags were programmed to sample GPS position (up to every minute), tri-axial acceleration (250 Hz), tri-axial magnetometer (50 Hz) and pressure (50 Hz). Mini sound tags also recorded audio data (96 kHz, 200 Hz – 48 kHz bandwidth). The active sonar within the sonar tags recorded acoustic backscatter returning from 10 µs pings with a centre frequency of 1.5 MHz at a 25 Hz ping rate. The active sonar operated with a 3.4° aperture beam width and a 6 m range (Goulet et al., 2019). Tags were set to record only during night hours (from 6 P.M. to 6 A.M. local time, i.e. foraging periods of AFS, (Boyd & Croxall, 1992)) to save battery. Tags were glued to the fur using quick-setting epoxy glue (Araldite AW 2101, Ciba) and recovered in January 2023 after a single foraging trip at sea using the same capture and sedation method.

### Replication statement

| Scale of inference | Scale at which the factor of interest is applied | Number of replicates at the appropriate scale |
| --- | --- | --- |
| Population | Population | 8 individuals |

### Data analyses

Data recovered from tags were analysed using custom-written codes and functions from www.animaltags.org in MATLAB version 2022b (The MathWorks, 2022). Statistical analyses were conducted in R software version 3.5.1 (R Core Team, 2018).

#### Prey capture attempt identification

Prey capture attempts (PrCAs hereafter) were detected from the 250 Hz tri-axial acceleration data recorded by the sonar tags, by computing the norm of the differential of the tri-axial acceleration (norm-jerk hereafter), as described in Chevallay et al. (2023). To establish a threshold for detecting prey strikes, the maximum norm-jerk value over consecutive 10 s intervals was log-transformed and displayed as histograms for each seal. These plots showed a bimodal distribution with a minimum at 3000 m.s^-2^ for all individuals, and was set as a threshold to identify strikes in the full norm-jerk series. As prey may be encountered in patches or may elude capture, leading to a bout structure in prey strikes, strikes occurring less than 15 s from the previous strike were grouped in the same PrCA bout (referred as bout hereafter) according to the distribution of inter-strike interval (Chevallay et al., in revision).

#### Sonar data analysis

Sonar data recorded during bouts were displayed as echograms, showing the time on the horizontal axis and the distance from the tag on the vertical axis, extending from 10 s before the bout start time to 2 s after the bout end. Different variables describing predator-prey interactions were extracted manually from echograms following the method described by Chevallay et al. (in revision): (1) number of prey, (2) prey evasive behaviour, (3) prey acoustic size and (4) echo intensity of the prey trace. (1) The number of prey were defined as the maximum number of independent echo traces within a same ping (Jones et al., 2008). It was scored as one, i.e. a single prey, two, or more than two, i.e. a school of prey. (2) Prey evasive behaviour was identified from the closing speed between predator and prey, which will vary in case of prey reaction, resulting in a change in the slope of the prey echo trace (Goulet et al., 2019; Vance et al., 2021). If a prey reaction was observed during the bout, prey was considered as evasive. (3) Prey acoustic size was estimated from the −20 dB echo pulse width measured on the widest part of the prey trace on evasive prey only (Burwen et al., 2003). (4) Echo intensity was defined as the maximum of the echo-to-noise ratio measured on the prey trace. Echo-to-noise ratio was computed as the subtraction between the intensity of the signal (in dB) and the noise level (in dB), defined as the 10^th^ percentile of the signal recorded in the last meter. It was measured only in prey traces visible in the 0.20-0.70 m range, to avoid measurement bias due to the distance between the target and the transducer. This range was chosen because it is where most of prey traces were visible.

#### AFS hunting behaviour

Metrics describing AFS fine-scale behaviour were extracted from the 250 Hz tri-axial acceleration data and the 50 Hz tri-axial magnetometer data. Posture of AFS was inferred from Euler angles (i.e. pitch angle (rotation around the left-right axis), roll angle (rotation around the longitudinal axis) and heading angle (rotation around the dorso-ventral axis, see Johnson and Tyack (2003) for details of the formulas). We also described adjustments in travel direction during the approach as a proxy for prey detection by computing the change in pointing angle, i.e. the angular change in direction of the longitudinal axis from the second before (Chevallay et al., 2023; Miller et al., 2004). Pointing angle was computed every second during the approach phase and we then computed the change in pointing angle as the temporal evolution of the pointing angle every second (Chevallay et al., 2023; Miller et al., 2004). Bouts were characterised in term of duration (time elapsed from the first and last strike of the bout), and intensity (RMS of the norm-jerk signal during the bout). Flipper strokes were detected from the dynamic acceleration of both the heave and the surge accelerometer axes (Jeanniard-du-Dot et al., 2016) by applying a high-pass filter with a cut-off frequency of 1.6 Hz, i.e. 70% of the dominant stroke frequency, on both axes. Absolute values of the dynamic heave and surge accelerations were then summed to obtain the swimming effort, a proxy of the AFS swimming activity (Aoki et al., 2011; Maresh et al., 2014).

### Statistical analyses

Prey vertical distribution was compared between prey types using generalized mixed models (GLMM hereafter, R package “MASS”, (Ripley et al., 2013)) with a gamma distribution, and with depth (m) as a response variable, prey type (schooling, single evasive or single non-evasive prey) as fixed effects, and individual seal identities as random effects. Acoustic size and echo intensity distribution of single and schooling prey traces were compared using Kolmogorov-Smirnov test. Approach behaviour between prey types were compared using GLMM with a gamma distribution, with bout duration (s), swimming effort (m.s^-2^), pitch, roll and heading extent (°) as response variables, prey type (schooling, single evasive or single non-evasive prey) as fixed effects, and individual seal identities as random effects. As the data represents a time-series, an autocorrelation structure of order 1 was added to the models. Results are all displayed as mean ± sd.

## Results

### Foraging behaviour

Tags were deployed during a single foraging trip at sea. Six AFS travelled South, on the edge of the Kerguelen plateau and two travelled East (Figure 1B).

**Figure 1:**
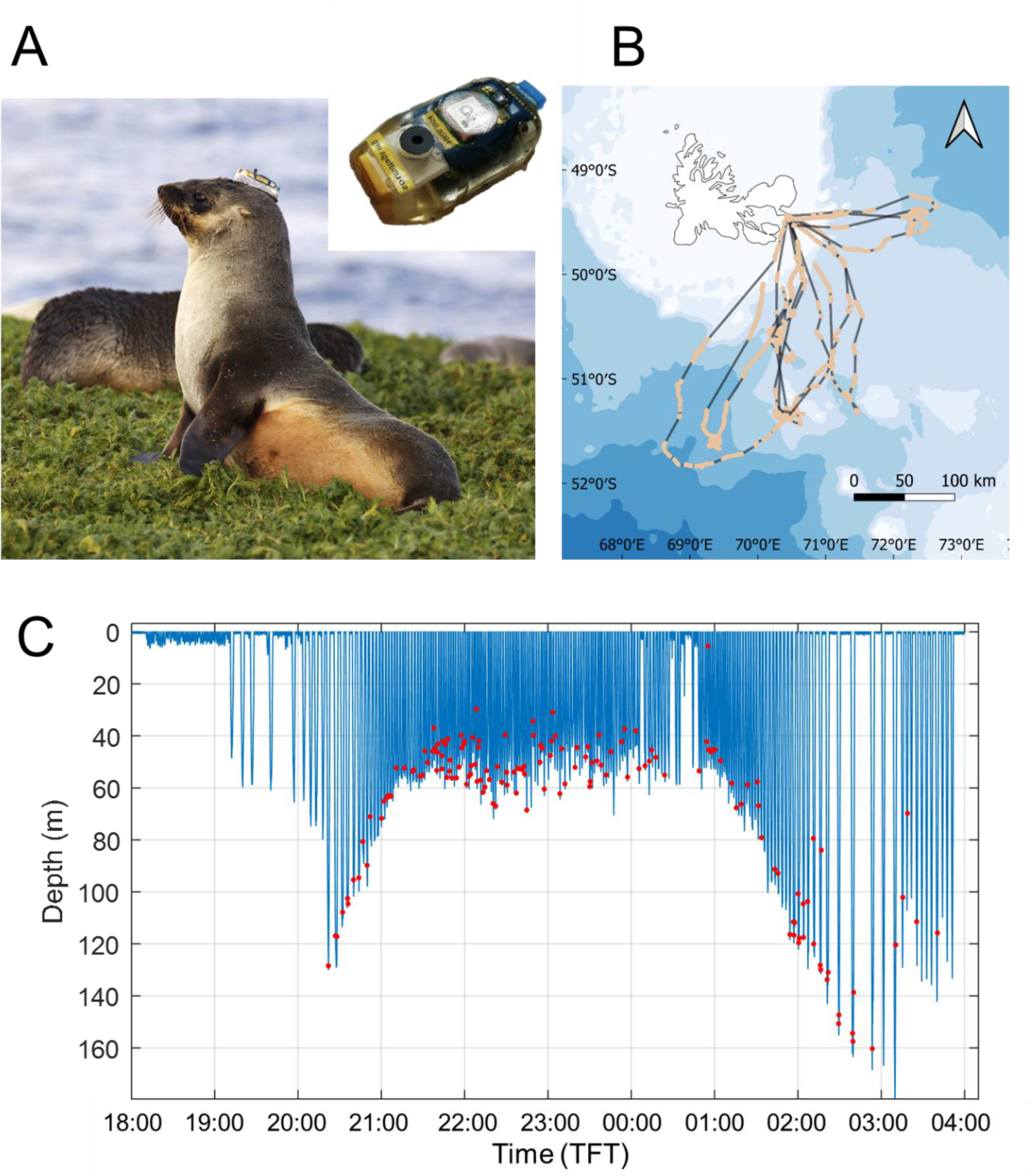
(A) Female AFS equipped with a sonar tag, pictured in the insert at the top right corner. (B) GPS track of eight female AFS from Pointe Suzanne colony on Kerguelen Islands equipped with sonar and sound tags in December 2022 during a single foraging trip at sea. Grey lines corresponds to night periods when the tag was switched off. (C) Example of a dive profile recorded during one foraging night for a female AFS. Red dots indicate PrCA bouts, identified as spikes in the norm of the differential of the tri-axial acceleration.

Foraging trips lasted 8 ± 1 d (min-max 6-10 d) per individual, and tags were set to record data only at night. During deployment, AFS performed 298 ± 129 dives per night (min-max 160-541) at 26.5 ± 27.8 m for 57.0 ± 53.6 s on average, with deeper dives performed at the beginning and the end of the night (Figure 1C). AFS performed 924 ± 165 bouts (min-max 622-1151 bouts) during the whole trip, i.e. 142 ± 20 bouts per night (min-max 104-166 bouts per night).

### Prey characteristics and distribution

Prey characteristics and behaviour were inferred on 1212 echograms with clear prey traces (Figure 2). Schooling prey represented 62% of targeted prey. Those prey usually formed large diffuse schools showing evasive behaviour (Figure 2C). Single prey represented 38% of targeted prey, among which 65% showed evasive behaviour (Figure 2A&B). Prey reacted 1.0 ± 2.3 s before the first strike of the bout, at a distance of 0.43 ± 0.21 cm from the AFS.

**Figure 2:**
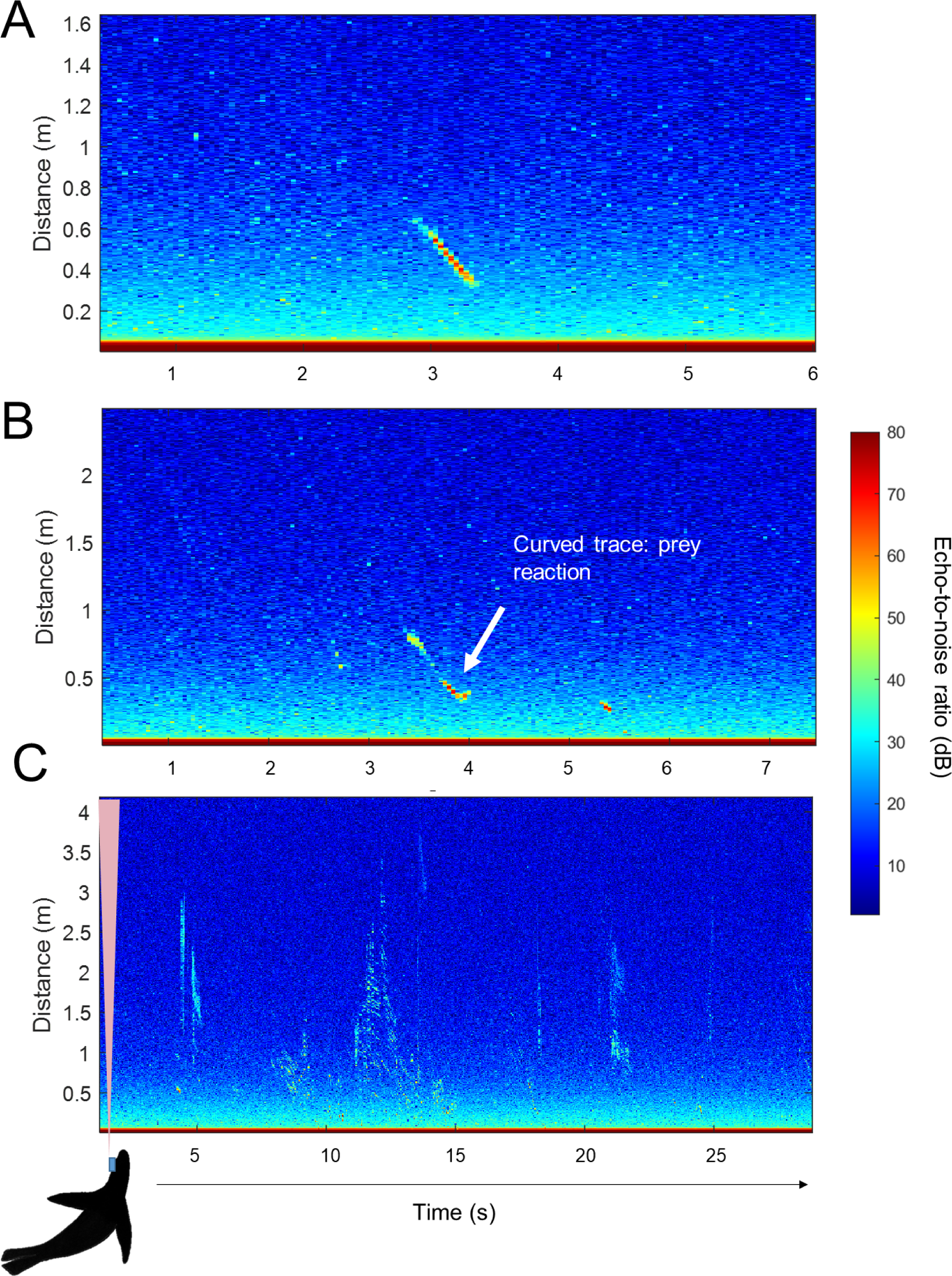
Echograms showing sonar data recorded during a PrCA bout by a female AFS equipped with a sonar tag in December 2022 at Pointe Suzanne, Kerguelen Island (time in s on the horizontal axis and distance from the sonar tag in m on the vertical axis). The colour scale indicates echo-to-noise ratio (ENR) on a dB scale. Echograms respectively display (A) a single non-evasive prey, (B) a single evasive prey and (C) schooling prey insonified over successive sonar pings. When independent echo traces were seen in the same ping, prey was considered as schooling prey. Prey evasive behaviour was identified from the closing speed between predator and prey, which will vary in case of prey reaction, resulting in a change in the slope of the prey echo trace.

Prey acoustic size ranged from 2.0 to 46.5 cm for single prey (6.3 ± 5.6 cm, Q1-Q3: 3.1-7.0 cm, Figure 3). Individuals prey within schools had significantly lower acoustic size than single prey, ranging from 1.0 to 7.4 cm (2.5 ± 1.1 cm, Q1-Q3: 1.6-3.1 cm, KS-test, P < 0.001, Figure 3). Echo intensity of single prey traces was significantly higher than of schooling prey traces (61.6 ± 9.2 dB, Q1-Q3: 55.9-67.7 dB for single prey; 50.3 ± 8.3 dB, Q1-Q3: 45.1-54.5 dB for schooling prey, KS-test, P < 0.001, Figure 3). Schooling prey were encountered slightly shallower than single prey (GLMM, P < 0.001, 42.1 ± 18.2 m vs. 49.3 ± 18.2 m respectively), while no difference in depth was found between single reactive and non-reactive prey (49.3 ± 23.0 vs. 48.2 ± 24.8 m, GLMM, P = 0.1603).

**Figure 3:**
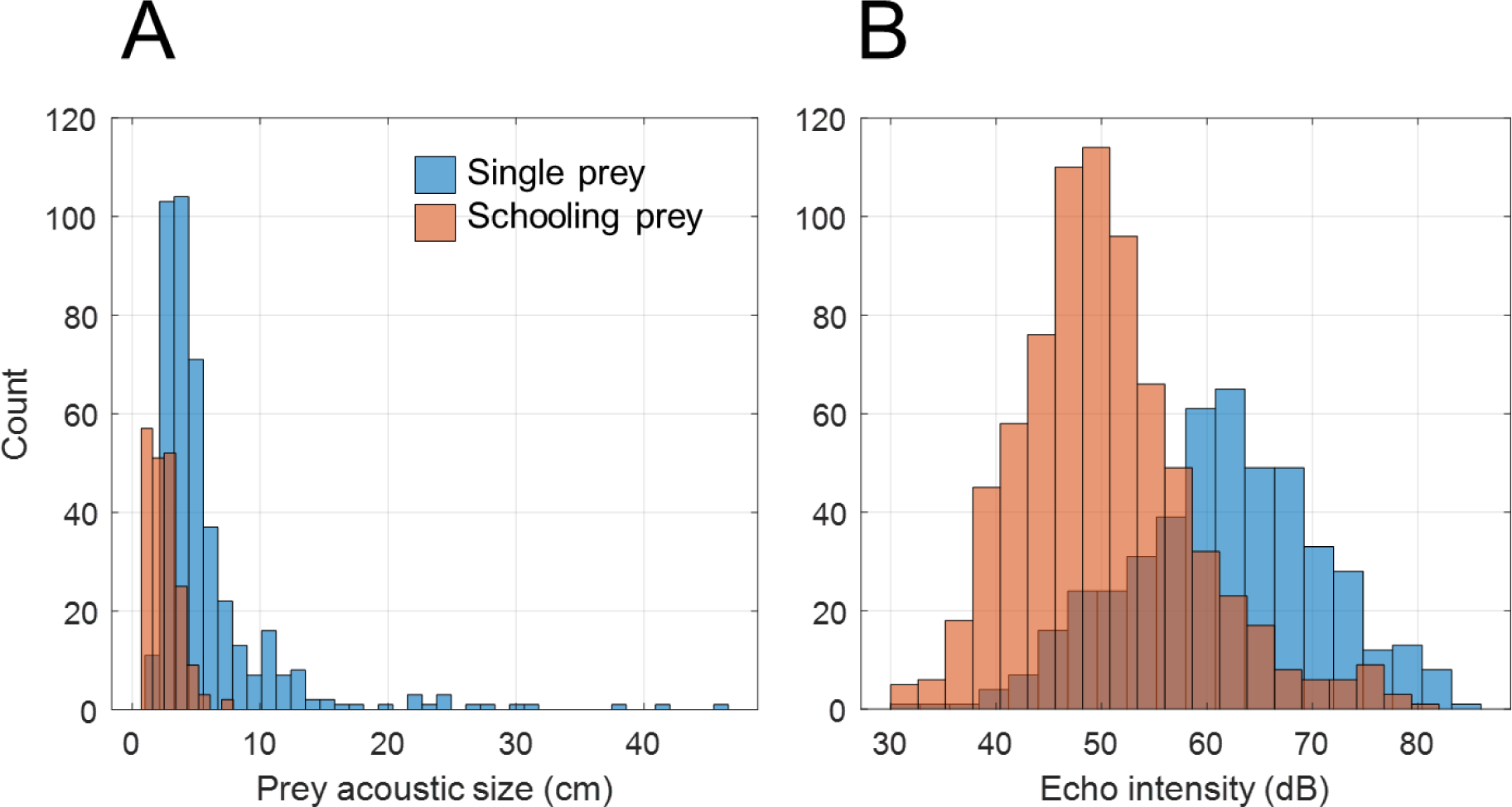
(A) Acoustic size and (B) echo intensity distributions of prey targeted by five female AFS equipped with sonar tags in December 2022 at Pointe Suzanne on Kerguelen Islands. Prey acoustic sizes were estimated on 419 single prey traces and on 199 schooling prey traces from the −20 dB echo pulse width measured on the widest part of the prey echo trace. Echo intensity was estimated on 467 single prey traces and 749 schooling prey traces and was defined as the maximum of the echo-to-noise ratio measured on the prey trace.

### Hunting behaviour

Between bouts, AFS generally glided resulting in a low swimming activity (Figure 4C), with AFS only giving 0.2 ± 0.1 flipper strokes per second, i.e. 1 flipper stroke every 5 s (Q1-Q3: 0.1 to 0.3 flipper strokes per second, i.e. 1 flipper stroke every 3 to 10 s). During the approach phase, i.e. 10 seconds before the bout start, swimming activity remained low, ranging between 0.1 and 0.6 flipper strokes per second (Q1-Q3), i.e. 1 flipper stroke every 1.6 to 10 s.

**Figure 4:**
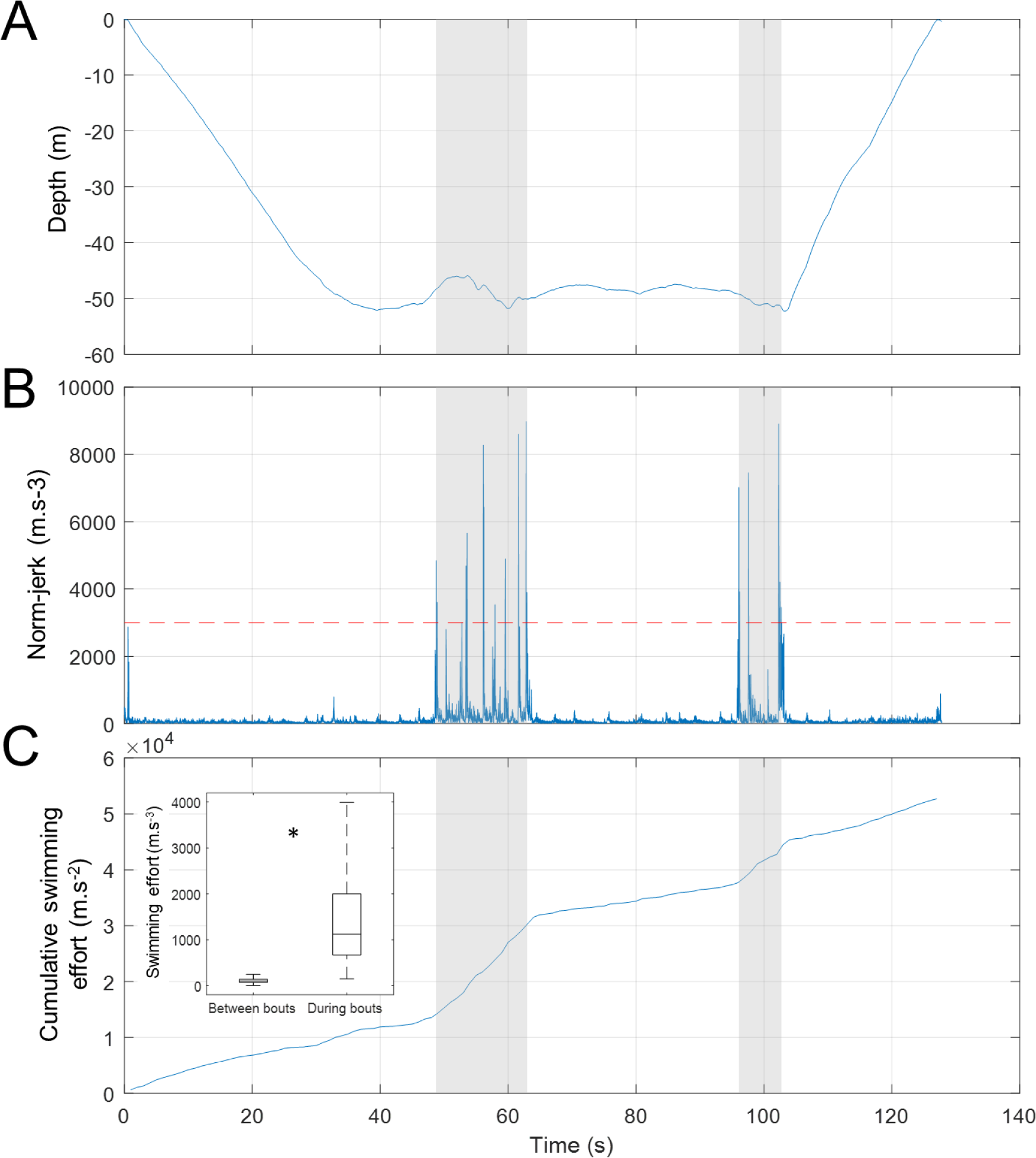
Example of a dive of a female AFS equipped with a sonar tag in December 2022 at Pointe Suzanne on Kerguelen Island. (A) Depth profile recorded during a single dive. (B) Norm of the differential of the tri-axial acceleration (norm-jerk) recorded during the dive. Spikes higher than 3000 m.s^-2^ were classified as PrCAs. Strikes occurring less than 15 s from the previous strike were grouped in the same bout, indicated by the grey shaded region. (C) Cumulative swimming effort, i.e. cumulative summed absolute values of the high-pass filtered surge and heave accelerometer axes, used as a proxy of the swimming activity. The insert represents the swimming effort per second between and during bouts. Asterisk indicates significant difference in swimming effort between and during bouts (GLMM, P < 0.001).

AFS usually adopted a horizontal posture (i.e. pitch angle close to 0°, Figure 5) during the approach and maintained this posture during the whole approach. For all individuals, roll angles were mainly positive and varied between 0 and 60°, translati ng into a right-tilted posture (Figure 5). AFS altered their direction of travel 1-2 s before the bout start (Figure 5).

**Figure 5:**
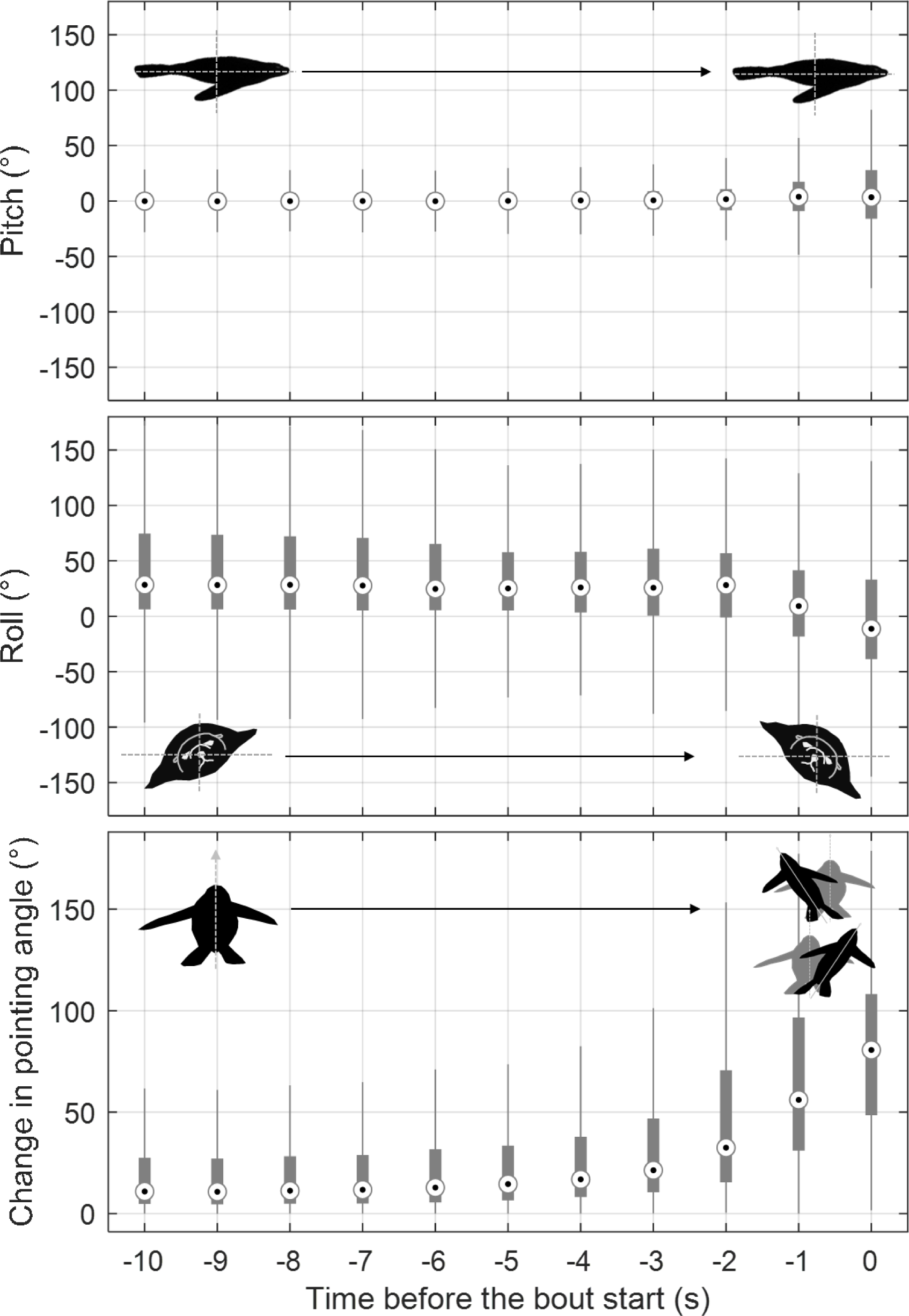
Posture of eight AFS equipped with sonar and sound tags in Kerguelen Islands in December 2022. Pitch and roll angles describe respectively the rotation around the left-right and longitudinal axes, while pointing angle reflects a change in direction of the longitudinal axis, likely associated with prey detection (Chevallay et al., 2023; Miller et al., 2004). Pointing angle was computed every second during the approach phase and we then computed the change in pointing angle relative to the previous second. They were computed on the static acceleration recorded by the tags during the approach phase, i.e. the 10 s preceding each bout (see Johnson and Tyack (2003) for details of the formulas).

AFS actively stroke flippers during bouts, resulting in a steady increase in swimming activity throughout bout duration (Figure 4). Swimming effort was significantly higher during bouts than during inter-bout periods (GLMM, P < 0.001, Figure 4). These bursts of acceleration lasted significantly longer for schooling prey and for single evasive prey than for single non-evasive prey (GLMM, P < 0.001, bout duration = 14.7 ± 12.6 s, 13.6 ± 10.3 s, and 7.2 ± 7.5 s respectively, Figure 6), with no difference between schooling prey and single evasive prey (GLMM, P = 0.7745). Similarly, bouts targeting schooling prey or single evasive prey were characterised by a higher number of head strikes (GLMM, P < 0.001, number of head strikes = 5.3 ± 4.6, 4.4 ± 2.5, 3.2 ± 2.0 respectively, Figure 6) with no difference between schooling prey and single evasive prey (GLMM, P = 0.3563). AFS swimming effort during the bout was significantly higher for schooling prey and single evasive prey than for single non-evasive prey (GLMM, P < 0.001, swimming effort = 14311 ± 12447 m.s^-2^, 12407 ± 8077 m.s^-2^, and 7258 ± 6270 m.s^-2^ respectively, Figure 6), but was not different between schooling prey and single evasive prey (GLMM, P = 0.1227). Finally, posture of AFS was highly variable during the bout regardless the type of targeted prey, with pitch, roll and heading extent being slightly inferior for single non-evasive prey, although not significantly (GLMM, P > 0.2834, Figure 6).

**Figure 6:**
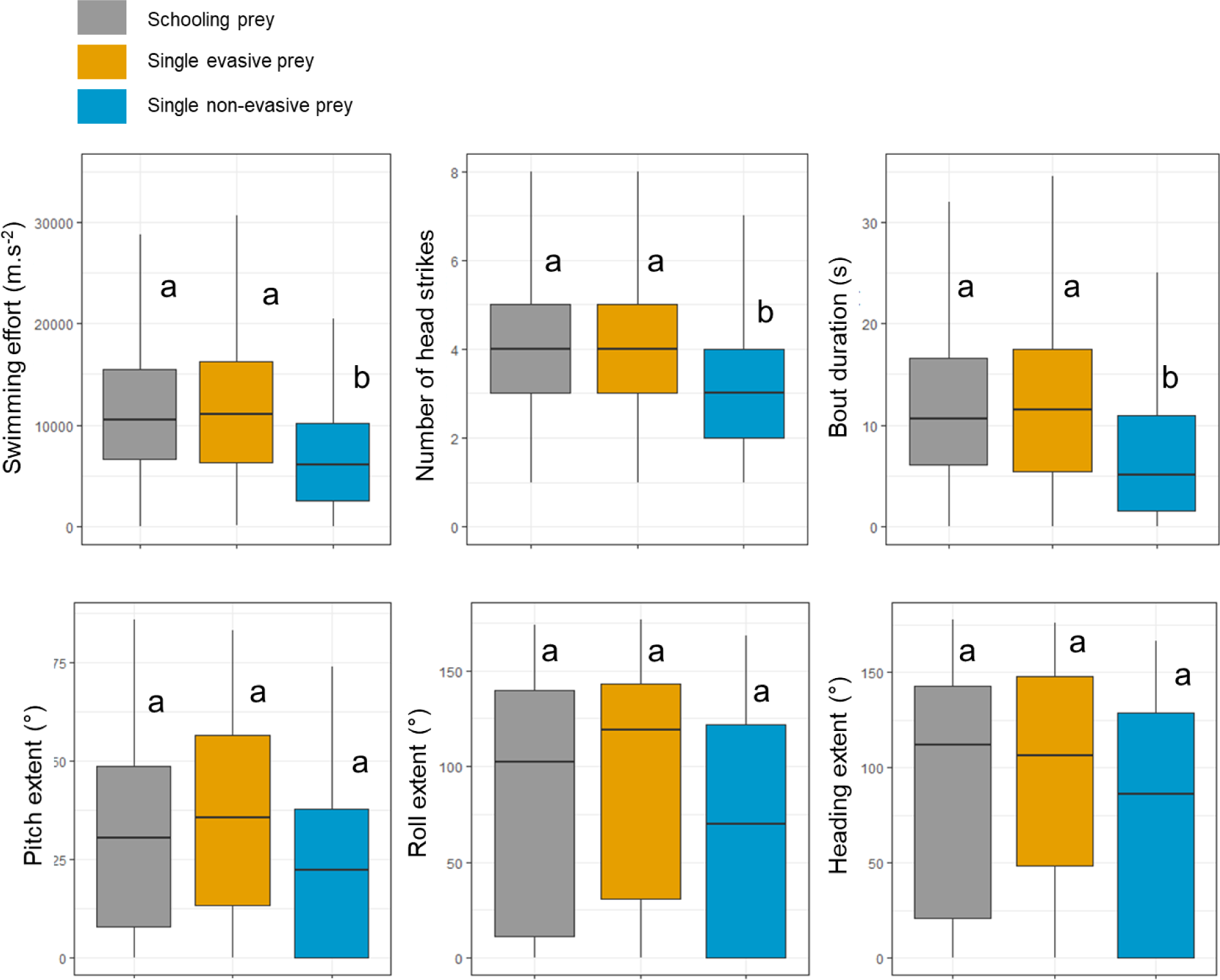
Hunting behaviour of five female AFS equipped with sonar tags at Pointe Suzanne on Kerguelen Islands in December 2022. Behavioural metrics were derived from accelerometer and magnetometer data recorded during bouts and were compared between prey types encountered by AFS. Prey types, i.e. schooling prey, single evasive prey, and single non-evasive prey were inferred visually from sonar data recorded during 1212 bouts. Letters in superscript indicate significant differences in behavioural parameters between prey types, i.e. schooling prey, single evasive prey and single non-evasive prey (GLMM, P < 0.05).

## Discussion

Inferring fine-scale characteristics of prey targeted by free-ranging diving predators and their detailed hunting tactics has been a long-standing technical challenge. New biologging devices combining ultra-high resolution movement sensors (i.e. accelerometers and magnetometers) with a synchronously-sampled high-ping rate active sonar had shed some lights on predator-prey interactions in deep-diving predators (Antoine et al., in revision; Chevallay et al., in revision; Goulet et al., 2019). Here and for the first time on otariids, we deployed sonar tags and high-resolution movement on AFS at Pointe Suzanne, Kerguelen Islands, to investigate prey selection and AFS hunting tactics.

### AFS prey characteristics and behaviour

The first objective of this study was to describe the fine-scale characteristics and behaviour of prey targeted by female AFS. Previous dietary work showed that female AFS from Kerguelen Islands mainly forage on myctophids, mostly from the genus *Gymnoscopelus* (Jeanniard-du-Dot, Trites, et al., 2017; Lea, Cherel, et al., 2002; Lea et al., 2008). Myctophids are small fish that are very abundant in the Southern Ocean, particularly around Kerguelen Islands. While they can be found at 1000 m depth during the day, most myctophid species perform diel vertical migration towards the surface at night. This makes them accessible to AFS that dive at ∼ 25 m depth on average (Boyd & Croxall, 1992). The main prey species targeted by females (i.e. *Gymnoscopelus piabilis*, *G. nicholsi* and *Electrona subaspera*) usually forms the upper-part of the mesopelagic community in the Polar Frontal Zone at night, migrating deeper than 300 m during daytime.

Size of prey found in the scats of female AFS was 4 to 15 cm (Lea, Cherel, et al., 2002), we estimated the acoustic size of prey targeted by tagged AFS between 3 and 7 cm. The smaller acoustic size of prey found in our study is not surprising. We have no control of the orientation of the prey regarding the sonar beam, and therefore the acoustic size can vary greatly depending on the orientation of the target relative to the beam. It is minimal when the target is perpendicular and maximal when oriented longitudinally to the beam. By measuring the maximum length of evasive prey only, we assumed that prey will be oriented longitudinally to the sonar beam at some point during the escape manoeuvre. This limits the target orientation bias and provides a reliable indication of prey actual size. In addition to the orientation, the composition, such as the presence of a gas-filled swimbladder in the target, can influence the measure of the acoustic size (Burwen et al., 2003). *Gymnoscopelus* species targeted by AFS do not have gas-filled swimbladder, responsible for most of the reflected acoustic signal (Dornan et al., 2019). Although the ultra-high frequency of the sonar tag makes it sensitive to targets with low acoustic reflexion properties (Goulet et al., 2019), the absence of gas-filled swimbladder might lead to an underestimation of the actual size of AFS prey. Even if there is still some uncertainty regarding the actual size of prey targeted by AFS using sonar recordings, this tag provides unique data on fine-scale prey characteristics during each prey encounter event which have been very difficult to obtain up to now.

Sonar recordings also allowed us to discriminate between single and schooling prey. We found that 62% of prey targeted by our tagged AFS were gathered in large diffuse and mobile schools encountered at shallower depths than single prey. Individual targets within schools seemed to have lower acoustic size compared to single prey items (i.e. 1-3 cm vs. 3-7 cm) and lower echo intensity (i.e. 45-54 dB vs. 56-68 dB). Differences in acoustic properties of single vs. schooling prey items could indicate different prey species (Misund, 1997). The large number of small individual targets within schools, and the evasive behaviour of schools could correspond to krill, which is known to form very mobile swarms (Hamner & Hamner, 2000). Recent trawl sampling and acoustic surveys performed on the Kerguelen plateau showed that one krill species, *Euphausia vallentini*, measuring 1-3 cm, is particularly abundant in the first 200 m of the water column (Cotté et al., 2022). *E. vallentini* is preyed upon by other Southern Ocean diving predators such as Macaroni penguins *(Eudyptes chrysolophus)*, rockhopper penguins *(Eudyptes chrysocome moseleyi)* or Gentoo penguins *(Pygoscelis papua)* (Ridoux, 1988; Tremblay & Cherel, 2003). AFS breeding in South Georgia are known to feed both on myctophids and Antarctic krill *(Euphausia superba)* (Reid & Arnould, 1996). It has been found that adult male AFS from Crozet Islands forage on *E. vallentini* (Cherel et al., 2009), however to our knowledge, this species has not been identified as a common prey species in the diet of female AFS from Kerguelen Islands (Jeanniard-du-Dot, Trites, et al., 2017; Lea, Cherel, et al., 2002; Lea et al., 2008). This could be artefactual, as crustacean remains are less conspicuous, more digestible and/or have a faster transit time more digestible than fish remains. DNA-metabarcoding analyses on scats should be conducted to verify the presence of krill in the diet of female AFS from Kerguelen (Jeanniard-du-Dot, Thomas, et al., 2017). We cannot also exclude a shift in the diet, as the last diet analyses were carried out twelve to twenty years ago.

### AFS fine-scale predator-prey interactions

The second objective of this study was to describe the AFS fine-scale behaviour during prey approaches and prey responses. We hypothesized that compared to Southern elephant seals, a phocid seal also feeding primarily on myctophids, AFS females adopt reactive fast-moving chases to target their prey. To test this hypothesis, we identified the timing of prey detection by AFS during their chase events. Using ultra-high resolution movement sensors (i.e. accelerometers and magnetometers), we identified the precise timing of prey strikes and then described the AFS direction of travel during the approach to find when they adjusted their course to intercept their prey (Chevallay et al., 2023; Miller et al., 2004). We found that AFS consistently changed their direction of travel 1-2 s before the strike. Swimming speed for AFS was estimated at less than 1 m/s during the bottom phase of the dive (Boyd et al., 1995), which gives a prey detection distance inferior to 1-2 m. Using sonar recordings, we were also able to identify prey reaction timing, estimated at approximately 1 s before the strike, i.e. at 0.4 m from the predator. Meanwhile, Southern elephant seals female adjust their direction of travel up to 10 s before the strike, i.e. 7-17 m prior the strike (Chevallay et al., 2023), with prey reacting 1 s before the strike, i.e. at 0.5 m from the predator (Chevallay et al. unpublished data). Therefore, our results suggest that, as opposed to elephant seals, AFS detect their prey very shortly before prey detects them, suggesting that they might be constrained to adopt a more reactive mode of hunting compared to elephant seals.

To detect prey, both elephant seals and AFS likely use a combination of whiskers and hearing (Adachi et al., 2022; Gläser et al., 2011; McCauley & Cato, 2016; Miersch et al., 2011). In particular, they might be able to locate large prey aggregations by listening to fish choruses (McCauley & Cato, 2016), as suggested in killer whales (*Orcinus orca,* (Guinet, 1992)), Indo-Pacific humpback dolphins (*Sousa chinsis*, (Barros et al., 2004)) or bottlenose dolphins (*Tursiops truncatus*, (Gannon et al., 2005)). However, tactile and acoustic cues may be difficult to detect in shallow dives due to turbulence and noise of waves. As AFS forage almost exclusively within the upper part of the mixed-layer depth, 20-40 m, their senses are likely affected by these wave movements. On the other hand, these turbulences might be attenuated under the mixed layer depth, the main foraging habitat of elephant seals, which forage at ∼ 200 to 600 m depths. These different environmental conditions may contribute to explain the shorter detection distance observed in AFS compared to SES. The smaller search swathe for AFS compared to SES suggest that they might need higher prey densities than SES to have the same probability of prey encounter as SES. Therefore, they likely need to concentrate their foraging activity on highly productive waters (Jeanniard-du-Dot & Guinet, 2021). AFS concentrate their foraging activity in the edge of the Kerguelen plateau, where the interactions between currents and continental slope enhance primary production and therefore promotes local aggregation of marine organisms (Lavoie et al., 2000; Meyer et al., 2015; Park et al., 2008).

A reactive mode of hunting requires the predator to perform fast manoeuvres to capture their evasive prey. We found that AFS rapidly increased their swimming effort and performed quick accelerations once interaction with prey begins. Bursts of accelerations lasted longer for evasive prey, likely reflecting prey pursuit. No matter the prey characteristics, we observed a high variability in AFS body angles, mostly in roll and heading angles, indicating turning and rolling manoeuvres during prey interactions. While AFS are highly mobile underwater, their small prey likely benefit from higher manoeuvrability (Domenici, 2001). However, their swimming speed is also much lower than that of AFS (i.e. 0.1-0.3 m/s for myctophids (Ignatyev, 1996) vs. cruising speed of 1-2 m/s for AFS (Boyd et al., 1995)) so it is likely that AFS have higher locomotor performances enabling them to catch up their prey quickly. The variability in AFS body angles during prey approaches seemed lower for single non-evasive prey than for single evasive prey, which could indicate that single non-evasive prey were approached in a straighter trajectory than evasive prey, and require less manoeuvres from AFS. This result was not significant, however our sample size was small and we observed a high inter-individual variability, which could explain why we did not detect any statistical differences in AFS posture between evasive and non-evasive prey.

Bursts of accelerations and active pursuit is also observed in other marine predators such as bottlenose dolphins (Maresh et al., 2004), short-finned pilot whales (*Globicephala macrorhynchus*, (Soto et al., 2008)), or sperm whales (*Physeter macrocephalus*, (Aoki et al., 2012)). Bursts of acceleration are highly energy-consuming especially for air-breathing diving predators that are limited by their oxygen stores. In sperm whales, bursts of acceleration can be responsible for up to 50% of energy consumption during dives (Aoki et al., 2012). In short-finned pilot whales, the energy spent in bursts of acceleration might explain their relatively short dive duration compared to diving predators of similar size (Soto et al., 2008). Similarly, AFS usually perform short dives of approximately 1-2 min (Boyd & Croxall, 1992), while smaller predators such as king penguins (*Aptenodytes patagonicus,* 10-15 kg vs. 30 kg for female AFS) or Adélie penguins (*Pygoscelis adeliae,* 3-8 kg) routinely dive during 3-5 min (Hanuise et al., 2013) and 1-2 min respectively (Chappel et al., 1993), even if these differences in dive capacities between species are also due to differences in physiological capacities (i.e. total oxygen stores, myoglobin concentration, etc.). Therefore, short dive durations observed in AFS might be explained by their active-hunting tactics. Indeed, estimations of energy expenditure using doubly labelled water techniques found that AFS had a field metabolic rate of ∼ 590 kJ.kg^-1^.day^-1^ (Jeanniard-du-Dot et al., 2016) while it was estimated at ∼ 100 kJ.kg^-1^.day^-1^ for Northern elephant seals (Maresh et al., 2014). These differences in energy expenditure might by partly explained by differences in hunting modes between these species, even if other factors such as thermoregulation capacities, diving capacities, oxygen stores might be taken into account to explain differences in pace of life between otariids and phocids (Jeanniard-du-Dot & Guinet, 2021; Ponganis, 2015). Active predators such as pilot whales or sperm whales usually select highly nutritious prey such as large muscular squids to compensate the energy costs associated with bursts of acceleration (Spitz et al., 2012). AFS mainly forage on myctophids, which are highly nutritious as they are rich in lipids (Lea, Nichols, et al., 2002). In particular, AFS target large species belonging to the genus *Gymnoscopelus* measuring up to 16 cm (Lea, Cherel, et al., 2002), so this active mode of hunting seems compensated by the consumption of large calorific prey.

Bouts associated with schooling prey were long with several successive head strikes (mostly between 3 and 5), high swimming efforts and high turning rates, suggesting that AFS might successively capture several prey within the school. We observed similarities in AFS behaviour during bouts associated with schooling prey and single evasive prey. Given the limited range of the sonar, it is likely that only tight aggregations appear as schools on echograms, while diffuse schools with a high inter-individual distance might be classified as single prey. Bouts associated with single evasive prey were significantly longer, associated with a higher swimming effort and a higher number of head strikes than bouts associated with single non-evasive prey, which might reflect successive attempts to capture evasive prey. However, long prey pursuit associated with prey that were classified as single evasive prey might also correspond to multi-prey engulfment within a scattered school. Similarly, we found a high variability in bout durations, number of head strikes and swimming efforts for all prey types including single non-evasive prey. While we would expect that these non-evasive prey would be captured in a single head strike without a chase, the longer bouts with several head strikes might correspond to multi-capture of non-mobile prey gathered in large scattered aggregations. A fine-scale analysis of AFS three-dimensional trajectory during bouts could help distinguishing between scattered schools and actual isolated prey.

While bursts of accelerations are associated with prey capture attempts, AFS generally displayed a low swimming activity between bouts giving only one flipper stroke every 3 to 10 s. Similar gliding behaviour was also observed in Southern elephant seals (Chevallay et al., 2023) and sperm whales (Aoki et al., 2012), in which gliding behaviour might help reduce the bow wave generated by their forward movement. Indeed, fish prey can detect turbulences produced by an approaching predator using their lateral lines. Consequently, AFS might decrease the risk of alerting their prey by lowering their swimming activity (Blaxter & Fuiman, 1990; McHenry et al., 2009; Stewart et al., 2013). Moreover, reducing body movements and the associated water turbulences might help AFS to better pick up tactile and acoustic cues of nearby prey in the turbulent surface waters of the windy Southern ocean, as suggested for rough-toothed dolphins (*Steno bredanensis,* (Götz et al., 2006)).

## Conclusion

This study provides insights into fine-scale prey characteristics and hunting tactics of free-ranging Antarctic fur seals. We found that AFS forage both on individual prey and on smaller prey items gathered in large diffuse and evasive schools, which may indicate a mixed diet between myctophids and krill. We suggest that AFS adopt a reactive mode of hunting relying on their ability to perform fast manoeuvres and bursts of acceleration. The active hunting tactics used by AFS is likely very energy-consuming but might be compensated by the consumption of highly nutritious prey. Here we compared the hunting tactics of AFS with another predator foraging on myctophids, the SES, and we showed that the two species adopt distinct hunting styles. As opposition to the active pursuit used by highly manoeuvrable AFS to capture their prey, low-expenditure swimming SES rely on higher detection capacities, allowing them to approach their prey stealthily without triggering an escape reaction and without spending too much energy. Differences in manoeuvrability, locomotor performances and detection capacities between AFS and SES might explain their differences in hunting styles.

## Acknowledgments

Field work in Kerguelen was supported by the French Polar Institute (Institut Polaire Français Paul Emile Victor) as part of the Ornithoeco programme (n. 109, PI C. Barbraud). We thank Nicolas Bonnetti, Lucas Bouland, Elie Castang, Pierre Guenot, Lola Gilbert, Camille Henriet, Ludovic Ivars and Sébastien Picon for their help in collecting the data. We want to thank Mark Johnson and Pauline Goulet for providing tags, software and codes for data analysis. We thank Benjamin Dupuis for his help in finding the title of this paper.

## Author contributions

MC, CG and TJDD conceived the ideas and designed methodology. MC and TJDD collected the data. MC and DGT analysed the data. MC, CG and TJDD led the writing of the manuscript. All authors contributed critically to the drafts and gave final approval for publication.

## Statement on inclusion

All authors were engaged early on with the research and study design to ensure that the diverse sets of perspectives they represent was considered from the onset. Whenever relevant, literature published by scientists from the region was cited.

## Data availability statement

Data used on the manuscript will be deposited on a Dryad depository at the time of publication.

